# Sensitivity to direction and velocity of fast frequency chirps in the inferior colliculus of awake rabbit

**DOI:** 10.1101/2022.10.26.513923

**Authors:** Paul W. Mitchell, Kenneth S. Henry, Laurel H. Carney

**Author notes:** Corresponding author Laurel H. Carney.

## Abstract

Neurons in the mammalian inferior colliculus (IC) are sensitive to the velocity (speed and direction) of fast frequency chirps contained in Schroeder-phase harmonic complexes (SCHR). However, IC neurons are also sensitive to stimulus periodicity, a prominent feature of SCHR stimuli. Here, to disentangle velocity sensitivity from periodicity tuning, we introduced a novel stimulus consisting of aperiodic random chirps. Extracellular, single-unit recordings were made in the IC of Dutch-belted rabbits in response to both SCHR and aperiodic chirps. Rate-velocity functions were constructed from aperiodic-chirp responses and compared to SCHR rate profiles, revealing interactions between stimulus periodicity and neural velocity sensitivity. A generalized linear model analysis demonstrated that periodicity tuning influences SCHR response rates more strongly than velocity sensitivity. Principal component analysis of rate-velocity functions revealed that neurons were more often sensitive to the direction of lower-velocity chirps and were less often sensitive to the direction of higher-velocity chirps. Overall, these results demonstrate that sensitivity to chirp velocity is common in the IC. Harmonic sounds with complex phase spectra, such as speech and music, contain chirps, and velocity sensitivity would shape IC responses to these sounds.

**Highlights:** IC neurons had diverse sensitivity to chirp velocity (speed and direction) of periodic and aperiodic-chirp stimuli.

Both velocity and periodicity sensitivity were necessary to predict neural responses to Schroeder-phase harmonic complexes.

Neurons were more commonly sensitive to the direction of chirps at lower-speeds (e.g., < 2 kHz/ms) than higher-speeds (e.g., > 2 kHz/ms) in the tested range.

The chirp speeds for which IC neurons were most sensitive are present in common harmonic sounds with realistic phase spectra, such as speech and music.

## 1. Introduction

The inferior colliculus (IC) receives convergent excitatory and inhibitory inputs from several auditory brainstem nuclei, and consequently displays a variety of complex response properties including frequency tuning (Davis, 2005), sensitivity to interaural differences (Yin et al., 2019), duration tuning (Casseday et al., 1994), and tuning to amplitude modulation (AM) (Langner and Schreiner, 1988; Krishna and Semple, 2000). Sensitivity of IC neurons to the velocity (speed and direction) of frequency sweeps has also been studied. Studies of bat IC responses report neurons that prefer, or even respond exclusively to, a particular sweep direction, a possible adaptation for echolocation (Suga, 1965; Fuzessery, 1994; Fuzessery and Hall, 1996; Gordon and O’Neill, 1998; Pollak et al., 2011). Additionally, sounds with statistics resembling vocal spectral-peak (formant) transitions have been used to produce IC spectrotemporal receptive fields (STRFs) in cats and bats (Escabí and Schreiner, 2002; Escabí et al., 2003; Andoni et al., 2007).

Recently, IC sensitivity to the velocity of sweeps has been explored in anesthetized gerbils (Steenken et al., 2022) and awake budgerigars (Henry et al., 2023). These studies used Schroeder-phase (SCHR) harmonic complexes, an advantageous stimulus for studying frequency-sweep sensitivity. The harmonic components of SCHR stimuli have flat magnitude spectra and curved phase spectra that create periodic acoustic waveforms with maximally flat temporal envelopes (Schroeder, 1970). The phase differences across harmonic components result in linear frequency sweeps within each fundamental period, extending from the lowest component, typically the fundamental frequency (F0) or its first harmonic, to the highest harmonic component.

As a tool for interrogating neural velocity sensitivity, the SCHR stimulus is unique in several ways. First, SCHR stimuli are notable for the speed of frequency sweeps, which travel from F0 to the maximum harmonic (here, 16 kHz) over the fundamental period (here, between 2.5 and 20 ms). The resulting velocities are faster than most commonly studied frequency-modulated signals, such as formant transitions (Liberman and Mattingly, 1989). We refer to these fast sweeps as SCHR chirps. Second, as a harmonic complex, the SCHR stimulus resembles voiced speech sounds. SCHR chirps emerge from phase differences between harmonic components. Similar phase differences are present in sounds like vowels, introduced into the glottal pulse by vocal-tract filtering. Therefore, neural sensitivity to chirp velocity would likely influence IC responses to speech sounds.

However, as harmonic stimuli, SCHR stimuli also have strong periodicity. IC neurons are notable for their tuning to the modulation frequency of amplitude-modulated (AM) sounds, often having enhanced or suppressed response rates to AM stimuli relative to an unmodulated stimulus (Krishna and Semple, 2000; Joris et al., 2004; Kim et al., 2020). It is unclear whether sensitivity to chirps requires a periodic context and whether the sensitivity is modulated by stimulus periodicity. Here, we use a new stimulus that isolated the velocity sensitivity from periodicity tuning. We recorded IC responses to both periodic SCHR stimuli and to aperiodic, random-chirp stimuli to assess velocity sensitivity. We explored the differences in prevalence of direction selectivity between the two stimuli and quantified the relative contributions of periodicity and velocity sensitivity in predicting SCHR response rates.

## 2. Methods

Extracellular, single-unit recordings were made in the central nucleus of the IC (ICC) of six female Dutch-belted rabbits (*Oryctolagus cuniculus*). Rabbits were obtained from Envigo (Denver, PA). The age of the animals during experiments ranged from 1-5 years, during which distortion-product otoacoustic emissions (DPOAEs) and IC neural thresholds were used to confirm normal hearing (Whitehead et al., 1992). Animals were housed in separate cages in the same room, with a 12-hour light cycle from 6 AM to 6 PM. Animals were removed from the study when their DPOAE magnitudes decreased or neural thresholds increased. All methods were approved by the University of Rochester Committee on Animal Resources.

### 2.1. Recordings

Recordings were made in daily two-hour sessions between 9 AM and 4 PM in sound-attenuated booths (Acoustic Systems, Austin, TX, USA). To prevent head movement of the awake rabbits, a headbar (metal or 3D-printed plastic) was surgically affixed to the skull using stainless-steel screws and dental acrylic. Rabbits were anesthetized using ketamine (66 mg/kg) and xylazine (2 mg/kg), administered intramuscularly, for headbar placement, craniotomy, and electrode placements (below). Rabbits were wrapped in a towel and secured in a custom-made chair to limit movement during recording sessions.

Recordings were made using four tetrodes, each consisting of four 18-μm platinum-iridium epoxy-coated wires (California Fine Wire Co., Grover Beach, CA, USA) and plated with platinum black to obtain impedances of approximately 0.1 – 0.5 MOhms. Tetrodes were surgically implanted in the ICC via a small (∼2-mm diameter) craniotomy in the top of the skull. Stainless-steel guide tubes protected the tetrodes, which were positioned just dorsal to the ICC and then advanced through the ICC using a manual microdrive (Five-drive, Neuralynx, Inc., Bozeman, MT, USA). Tetrodes were advanced before or after recording sessions. The microdrive and tetrodes were surgically replaced every few months, as needed, to vary the recording location. Neural recordings were made using an RHD recording system (Intan Technologies, LLC., Los Angeles, CA, USA), with Intan software running on a Windows PC.

Stimuli were generated using custom MATLAB code (Mathworks, Natick, MA, USA), a MOTU audio interface (16A, Mark of the Unicorn, Cambridge, MA, USA), and a Benchmark digital-to-analog converter and amplifier (DAC3 HGC, Benchmark Media Systems, Inc., Syracuse, NY, USA). Etymotic ER2 earphones (Etymotic Research, Inc., Elk Grove Village, IL, USA) were used to present sound stimuli via custom ear molds (Hal-Hen Company, Inc., Garden City Park, New York, USA). Prior to every session, stimuli were calibrated using ER-7C probe-tube or ER-10B+ microphones (Etymotic Research, Inc., USA). The calibration curve (magnitude and phase) was included in a pre-emphasis filter that compensated all stimuli for the frequency response of the acoustic system.

Action potential times and waveforms were extracted from neural recordings using custom MATLAB applications. To identify single-unit events, the voltage recording was filtered using a 4^th^-order Butterworth bandpass filter (300 – 3000 Hz). An action potential was identified when the filtered voltage signal exceeded a threshold, defined as four standard deviations of the voltage signal. Action potentials were sorted into clusters, presumed to represent different neurons, using features of the waveforms, primarily the slope of the waveform repolarization (Schwarz et al., 2012). Single-units were identified when less than 2% of the interspike intervals were shorter than 1 ms. Neurons studied in consecutive sessions were considered to be unique only when both tetrode location and response properties changed.

### 2.2. Stimuli

A set of stimuli was presented to define neural response properties, including frequency response maps, modulation transfer functions, SCHR sensitivity, and rate-velocity functions for aperiodic-chirp stimuli. To generate frequency response maps, a randomly-ordered sequence of 0.2-s-duration tones at frequencies ranging from 250 Hz – 16 kHz was presented at 13, 33, 53, and 73 dB SPL. Tones were presented either contralaterally or diotically with 10-ms raised-cosine on/off ramps, and were separated by 0.4 s of silence. Each stimulus condition was presented three times. Response maps, plots of average discharge rate vs. frequency at each stimulus level, were used to estimate characteristic frequency (CF), the frequency for which threshold was lowest.

Modulation transfer functions (MTFs) were estimated by computing the average rate in response to 100% sinusoidally amplitude-modulated noise as a function of modulation frequency, over a range of 2 – 350 Hz, with 3 steps per octave. The wideband noise (100 Hz – 10 kHz) was 1 s in duration, including 50-ms raised-cosine on/off ramps, presented diotically. Modulation frequencies were presented in random order, with 5 repetitions of each. Noise presentations were separated by 0.5 s of silence. Stimuli were presented at a spectrum level of 33 dB (overall level of 73 dB SPL). MTF categorization was based on criteria that were slightly modified from Kim et al. (2020). MTF types were categorized as band-enhanced (BE) if (1) the MTF had two consecutive points significantly greater than 120% of the unmodulated rate (*r*_*um*_), and (2) the MTF did not have more than one point significantly below 80% of *r*_*um*_. Similarly, MTFs were categorized as band-suppressed (BS) if (1) the MTF had two consecutive points significantly less than 80% of *r*_*um*_, and (2) the MTF did not have more than one point significantly above 120% of *r*_*um*_. MTFs that had both enhanced and suppressed bands that met the above criteria were categorized as hybrid MTFs. In the population of MTFs studied (N = 335), 177 (52.8%) were BS, 73 (21.8%) were BE, 51 (15.2%) were hybrid, and 34 (10.2%) were flat.

SCHR stimuli were produced using the phase spectrum introduced in Schroeder (1970) (Fig. 1A-D). For a harmonic complex with *n* components, the phase of the *n*th component was described by *θ*_*n*_ = *Cn*(*n* + 1)/*N*, where *N* is the total number of harmonic components, and *C* is a scalar that describes the phase-direction and duty cycle of the stimulus. SCHR stimuli with a positive C-value will be referred to as downward SCHR, reflecting the direction of their frequency chirps. SCHR stimuli with a negative C-value will be referred to as upward SCHR. Note that the direction of SCHR chirp (upward or downward) is the opposite of its phase (negative or positive, respectively). A random sequence of SCHR stimuli was presented, with F0 equal to 50, 100, 200, or 400 Hz, and C equal to -1, -0.5, 0.5, or 1. For all cases, the frequency of the highest harmonic component was 16 kHz. The number of harmonic components, *N*, depended on F0, and ranged between 40 and 320. Thirty repetitions of each F0 and C combination were presented. Stimuli were presented diotically at 63 dB SPL, with a duration of 0.4 seconds, 0.6 seconds of silence between stimuli, and 25-ms raised-cosine on/off ramps. Average discharge rates over the stimulus duration were calculated for each F0 and C combination, excluding a 25-ms window after stimulus onset. Receiver-operating-characteristic (ROC) analysis (Egan, 1975) was used to evaluate neural direction selectivity for SCHR conditions with equivalent F0 and C-values of opposite signs (e.g., F0 = 50 Hz, C = ±1). For this analysis, the response rates for each stimulus repetition were gathered, for chirps of equal speed but opposite direction. Then, ROC was used to measure the discriminability of chirp direction based on response rates of single stimulus repetitions. Neurons were deemed significantly biased toward one sweep direction if the ROC area under the curve (AUC) was above 70.7% or below 29.3%. The two separate ROC criteria were used to differentiate between selectivity for upward and downward chirps (akin to a two-tailed t-test). Neurons with AUCs above 70.7% were designated as selective for upward chirps (indicating greater mean response rate in response to upward chirps) and neurons with AUCs below 29.3% were designated as selective for downward chirps. The ROC criteria of 70.7% and 29.3% were chosen because they are equivalent to the targeted threshold of a two-down, one-up psychophysical procedure (Levitt, 1971).

**Figure 1.**
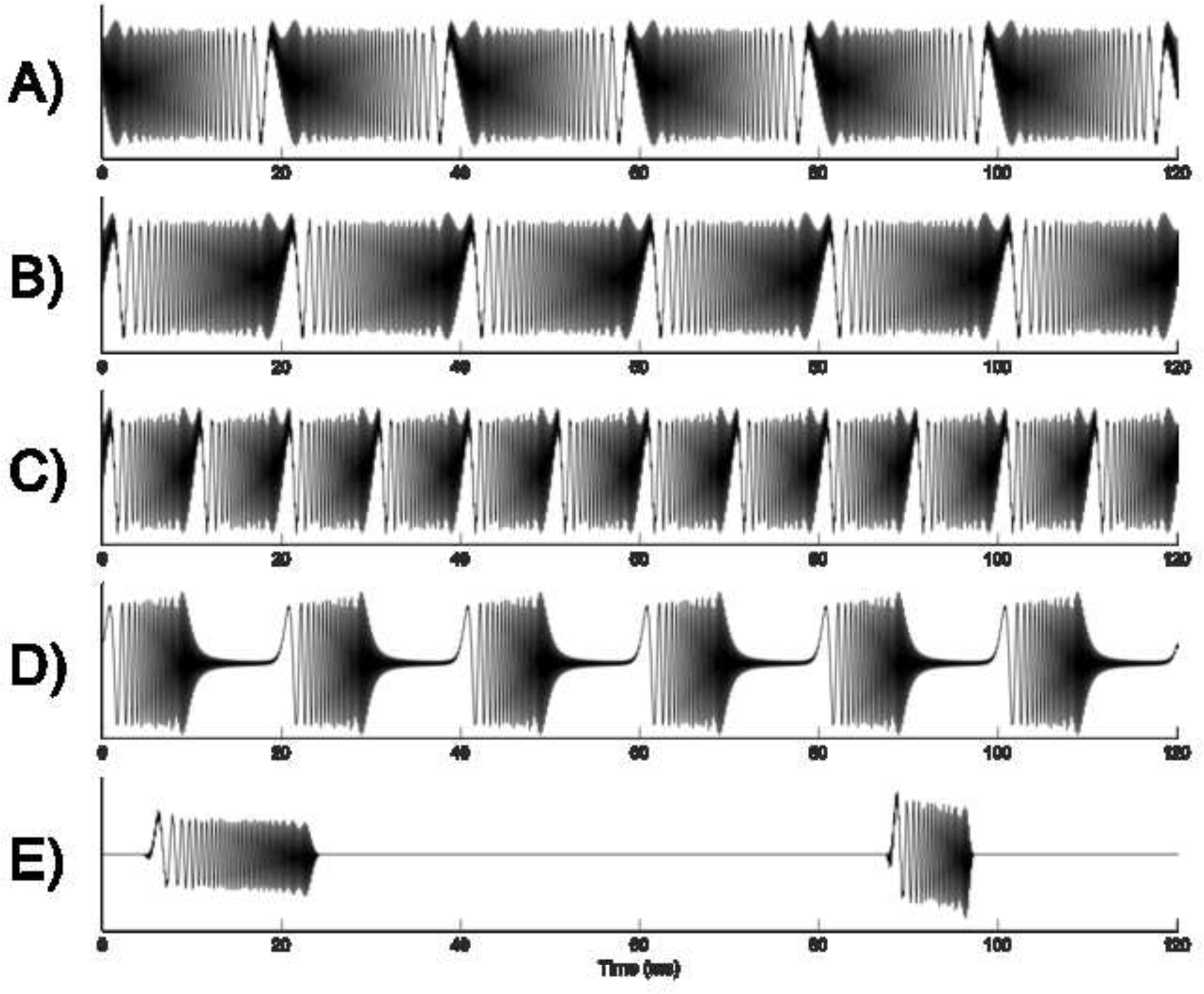
Stimulus waveforms of SCHR and aperiodic-chirp stimuli. Note that the maximum harmonic in this figure is 5 kHz for clarity; in the actual stimuli, maximum harmonic was 16 kHz. (A) Downward SCHR with F0 = 50 Hz, C = +1. (B) Upward SCHR with F0 = 50 Hz, C = -1. (C) Upward SCHR with F0 = 100 Hz, C = -1. (D) Upward SCHR with F0 = 50 Hz, C = -0.5. (E) Two consecutive aperiodic chirps separated by silence.

Rate-velocity functions (RVFs) were based on responses to aperiodic-chirp stimuli consisting of isolated SCHR chirps separated by silent periods of variable duration. These chirps were single fundamental periods extracted from a SCHR stimulus, equivalent to one full SCHR chirp spanning F0 to the maximum harmonic, or vice versa (Fig. 1E). To produce the stimulus waveforms, first, we generated a random sequence of chirp velocities that were equivalent to SCHR F0s of 25, 50, 100, 200, 400, and 600 Hz. Chirp direction (upward or downward, i.e., C = -1 and C = 1, respectively) was also randomly varied across stimuli. These parameters together resulted in aperiodic chirps with velocities ±0.40, ±0.80, ±1.59, ±3.16, ±6.24, and ±9.24 kHz/ms. Each combination of velocity and direction was presented a total of 42 times, in random order. To avoid periodicity, chirps were randomly spaced, using 40 – 60 ms uniformly distributed inter-chirp intervals. The full stimulus set was presented 20 times. Each individual chirp had raised-cosine on/off ramps, each with duration equivalent to 10% of chirp duration. Chirps were presented diotically at a sound level equal to 68 dB SPL − 10 × log_10_(*T*/*T*_*ref*_), where *T* is the duration of the chirp, and *T*_*ref*_ = 2.5 ms (the duration of a SCHR chirp with F0 = 400 Hz). This scaling ensured that energy was equivalent across chirps of different durations.

Response rate was calculated by summing spikes over a 15-ms time window starting at an estimate of the neural latency (Fig. 2) based on the response to a 73-dB tone at CF (from the response map). In cases where a neuron responded to 73-dB SPL tones over a wide frequency range, the neural latency to a chirp was sometimes substantially shorter than would be predicted by the latency at CF, as expected if instantaneous frequencies well below or above CF excited the neuron. In these cases (approximately a quarter of the neurons), the latency estimates were adjusted based on the frequency range over which the neuron responded. ROC analysis of aperiodic-chirp rates was performed in an identical manner as for SCHR rates. Aperiodic-chirp rates were used to generate an RVF, which described how a neuron’s rate changed as a function of chirp velocity (Figs. 3-4 below).

**Figure 2.**
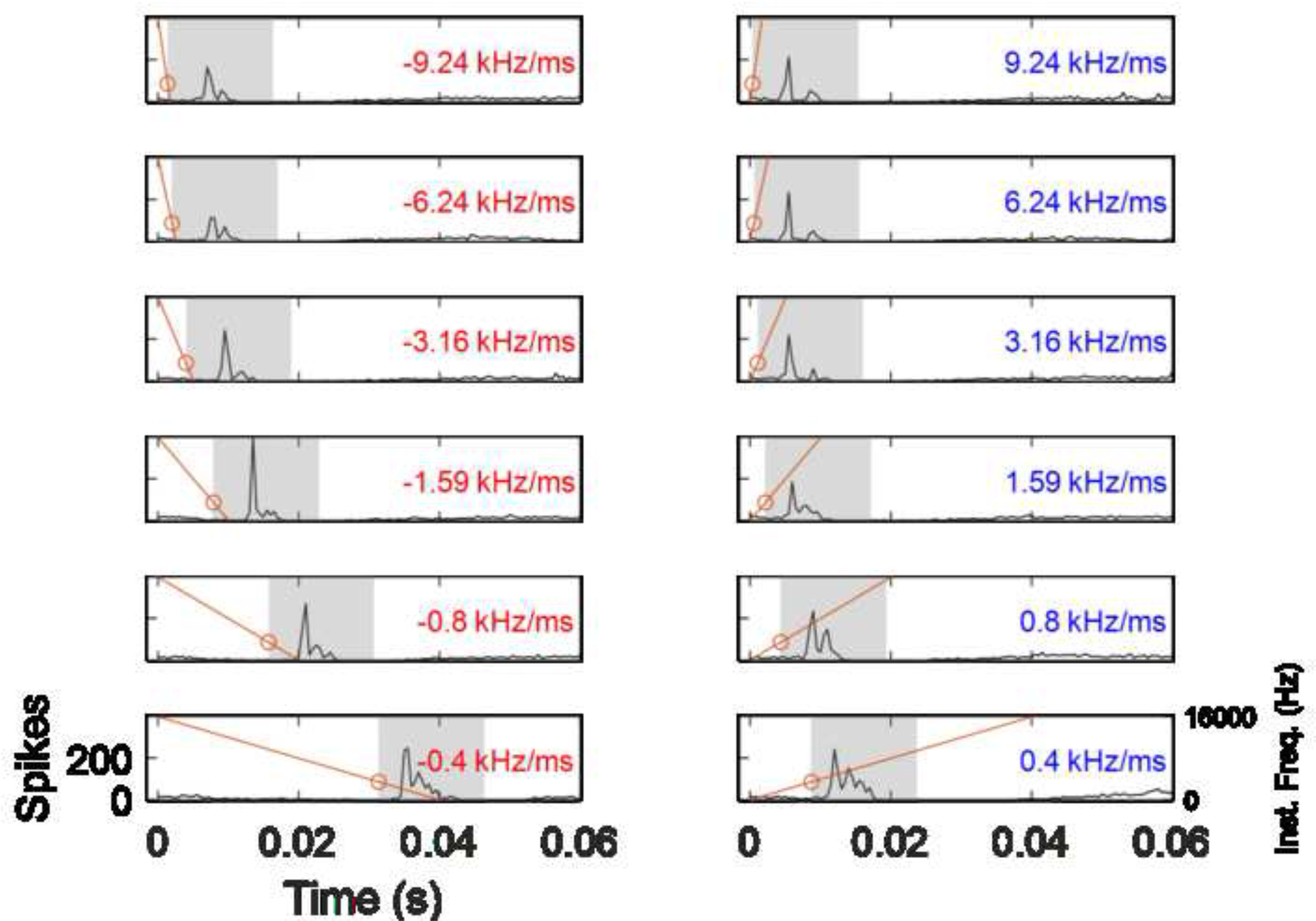
A visual depiction of aperiodic-chirp stimulus rate calculation. Black lines illustrate PSTHs (bin size = 0.5 ms) of responses to chirps of different velocities. Orange lines indicate the instantaneous frequency of the aperiodic chirp over time. The left column (red text) shows downward chirps, and the right column (blue text) shows upward chirps. Orange circles indicate when the instantaneous frequency of the chirp passes CF (here, 3500 Hz). The shaded rectangles indicate the placement of the 15-ms window over which response rate was calculated.

### 2.3. Analysis

Principal Component Analysis (PCA) was performed to identify the salient features in RVFs, using the MATLAB function PCA (2021b, MathWorks). PCA analysis allowed grouping of RVFs by their shape to gain insight into underlying patterns or trends. RVFs were normalized by their individual peak rates before PCA analysis. SCHR responses were predicted based on MTFs and RVFs using generalized linear models (GLMs) (Nelder and Wedderburn, 1972). First, SCHR rates, MTFs, and RVFs were normalized by each of their maximum rates. For the responses to periodic stimuli, (MTF and SCHR), cycle rate was used, defined as the average rate per fundamental period. Cycle rate (spikes/cycle) was necessary to compare responses to periodic stimuli and aperiodic chirps. For the aperiodic-chirp stimulus, rate was calculated using the method described above (Fig. 2). The periodicity and velocity of each SCHR stimulus was matched with the closest available modulation frequency and velocity from the MTF and RVF, respectively (data were not interpolated), creating a feature vector of the responses to periodicity and velocity changes. Then, a relationship between the periodicity and velocity feature vectors and SCHR rates was defined for the GLM, as follows:

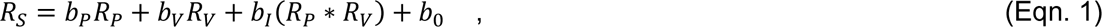

where *R*_*P*_ and *R*_*V*_ refer to rates of the periodicity and velocity feature vectors, respectively. *R*_*S*_ is the predicted response to SCHR stimuli in cycle rate, and *b*_*n*_ are the GLM coefficients. There were three terms in the GLM—a periodicity term, a velocity term, and an interaction term. GLM coefficients were estimated using the MATLAB function glmfit (2021b, MathWorks), with the distribution argument set to *normal* and constant argument set to *on*. The GLM coefficients were *b*_*P*_, the periodicity coefficient, *b*_*V*_, the velocity coefficient, *b*_*I*_, the interaction-term coefficient, and *b*_0_, a constant. A two-sided t-test was performed on the coefficients against the null hypothesis that the value of the coefficient was zero. After the SCHR rate prediction was calculated using estimated coefficients, the coefficient of determination (*R*^2^) of the predicted and recorded SCHR cycle rates was determined to assess prediction quality. This process was repeated for two additional, simpler GLMs:

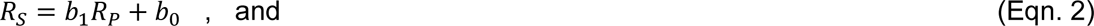

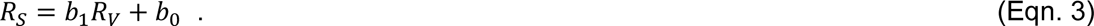

These simpler GLMs depended only on one feature vector and thus could be used to assess the importance of periodicity or velocity alone. The coefficients of determination (*R*^2^) of these two model predictions were obtained and used to measure the impact of the two features on neurons’ SCHR responses.

The following equation was used to calculate the contribution of velocity to the total explainable variance:

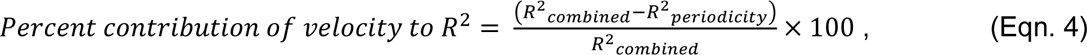

where *R*^2^_*combined*_ and *R*^2^_*periodicity*_ refer to the coefficient of determination of the combined model (Eqn. 1) and periodicity-alone model (Eqn. 2), respectively. Equation 4 allowed the amount of additional variance explained by the inclusion of velocity (either in the velocity term or interaction term in Eqn. 1) to be directly quantified.

## 3. Results

### 3.1. Examples of responses to SCHR and aperiodic-chirp stimuli

Data were recorded over a span of 330 sessions and included recordings from 335 unique single units. Response properties of single-unit neurons are illustrated using frequency response maps, MTFs, SCHR rate functions, and RVFs (Figs. 3 and 4). Frequency response maps were used to determine CF; for neurons with spontaneous activity, these plots also revealed off-CF inhibitory regions. MTFs were categorized by shape (i.e., BE, BS, hybrid, or flat). SCHR rate functions were generated by summing the total response over the duration of the periodic SCHR stimulus. Lastly, RVFs, which reflect velocity sensitivity in the absence of a periodic context, were constructed from aperiodic-chirp responses. Altogether, these stimulus responses were used to assess the interactions of different feature sensitivities, particularly to periodicity and velocity, in determining the SCHR responses.

**Figure 3.**
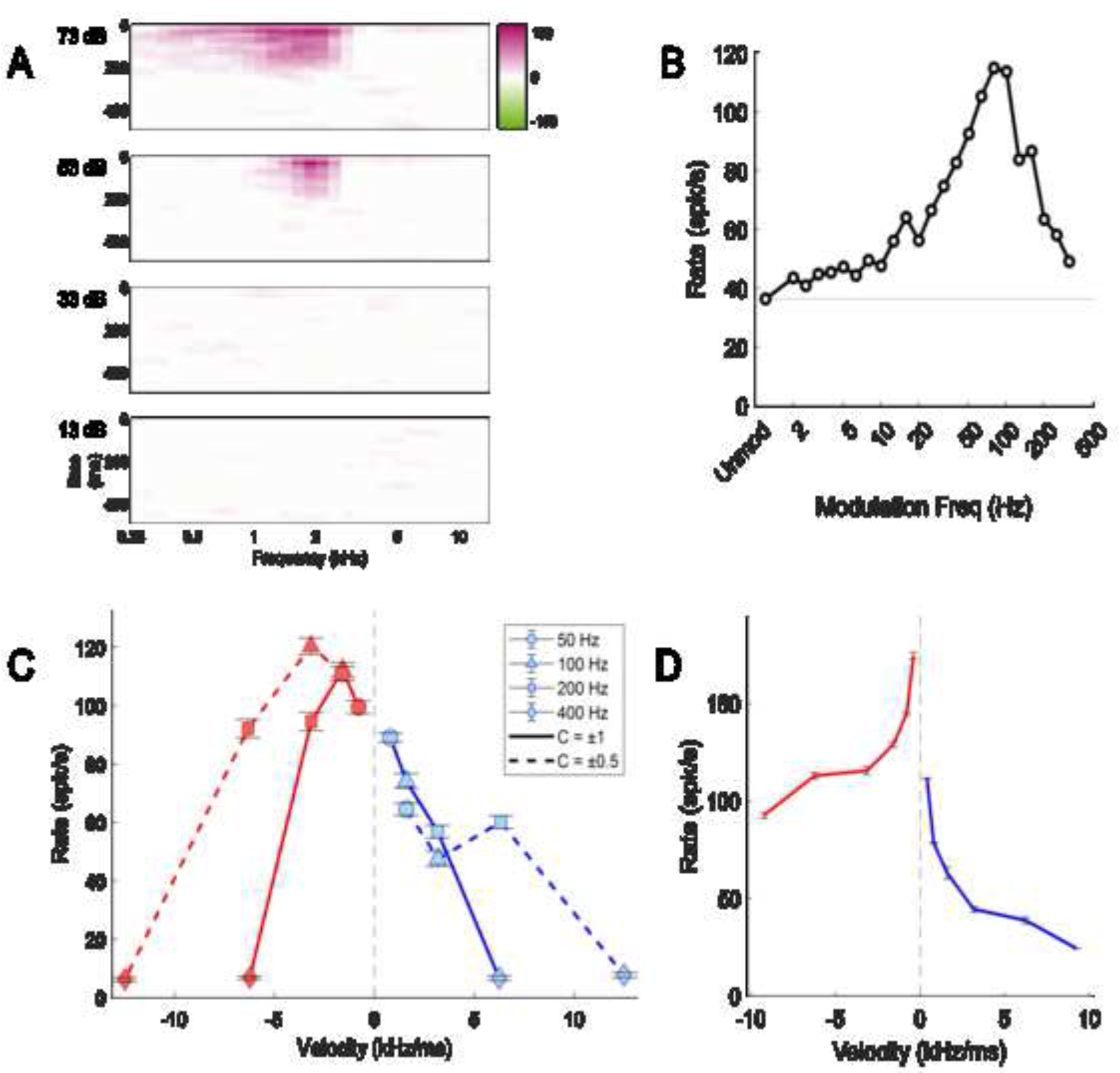
Responses of unit R24TT2P8N4 to characterizing stimuli. (A) Response Map – Panels are arranged vertically by sound level of pure tones. Color plots reflect response rate versus frequency (x-axis) and time following tone onset (y-axis); magenta regions have increased activity relative to spontaneous rate, and green regions have decreased activity relative to spontaneous rate. (B) Modulation Transfer Function – rate versus modulation frequency. The grey horizontal line reflects unmodulated rate. (C) SCHR response rates plotted as a function of chirp velocity. Solid lines are used for C = ±1, and dashed lines are used for C = ±0.5. Line color denotes direction of chirps (red—downward chirp, blue—upward chirp). Symbols are used to reflect SCHR F0, with circles, triangles, squares, and diamonds corresponding to 50, 100, 200, and 400 Hz respectively. (D) Rate-velocity function – constructed using responses to aperiodic-chirp stimuli. Error bars are the standard error of rates across repetitions (20 each).

Example responses for two neurons are illustrated in Figs. 3 and 4. The neuron in Fig. 3 had a BE MTF (Fig. 3B) and a predominantly excitatory response map with a CF of 2 kHz (Fig. 3A). This neuron had chirp sensitivity in response to SCHR stimuli that was influenced by stimulus periodicity. In the SCHR rate function, for C = ±1 (Fig. 3C, solid lines) a greater response rate to downward chirps was observed for all but the highest velocity. This velocity corresponded to a 400-Hz F0, a modulation rate for which the neuron did not have an enhanced rate in its MTF. In comparison, the RVF (Fig. 3D) had a stronger rate selectivity for downward chirps at all velocities, including the high-velocity range at which the SCHR rates were not selective. Thus, the neuron’s sensitivity to the velocity of SCHR chirps was impacted by stimulus periodicity, albeit both the SCHR responses and RVF were generally selective for the same chirp direction. It is notable that the C = ±0.5 SCHR responses (Fig. 3C, dashed lines) also differed from the responses to the aperiodic RVF, with rates that tapered off as modulation rate increased.

Other neurons’ SCHR responses were more strongly impacted by their periodicity tuning, even showing opposite chirp direction selectivity to that of their RVF. An example of such a neuron is shown in Fig. 4 which had a BS MTF (Fig. 4B) with the suppressed band approaching zero rate. The response map (Fig. 4A) was tuned to 1.2 kHz with a low threshold (13 dB SPL), and a high-frequency inhibitory region at higher sound levels. This unit’s SCHR response (Fig. 4C) and RVF (Fig. 4D) were quite different from one another. The SCHR rate function had higher responses to downward chirps at most velocities. In contrast, the RVF displayed a higher rate for upward chirps for most velocity pairs. Notably, for C = ±1 SCHR stimuli (Fig. 4C, solid lines), the two lowest velocity-pairs corresponded to modulation rates in the trough region of the BS MTF. While the mechanism for this apparent direction reversal cannot be ascertained from the MTF alone, it is apparent that the periodic nature of the SCHR signal can change the chirp-direction sensitivity for some neurons.

**Figure 4.**
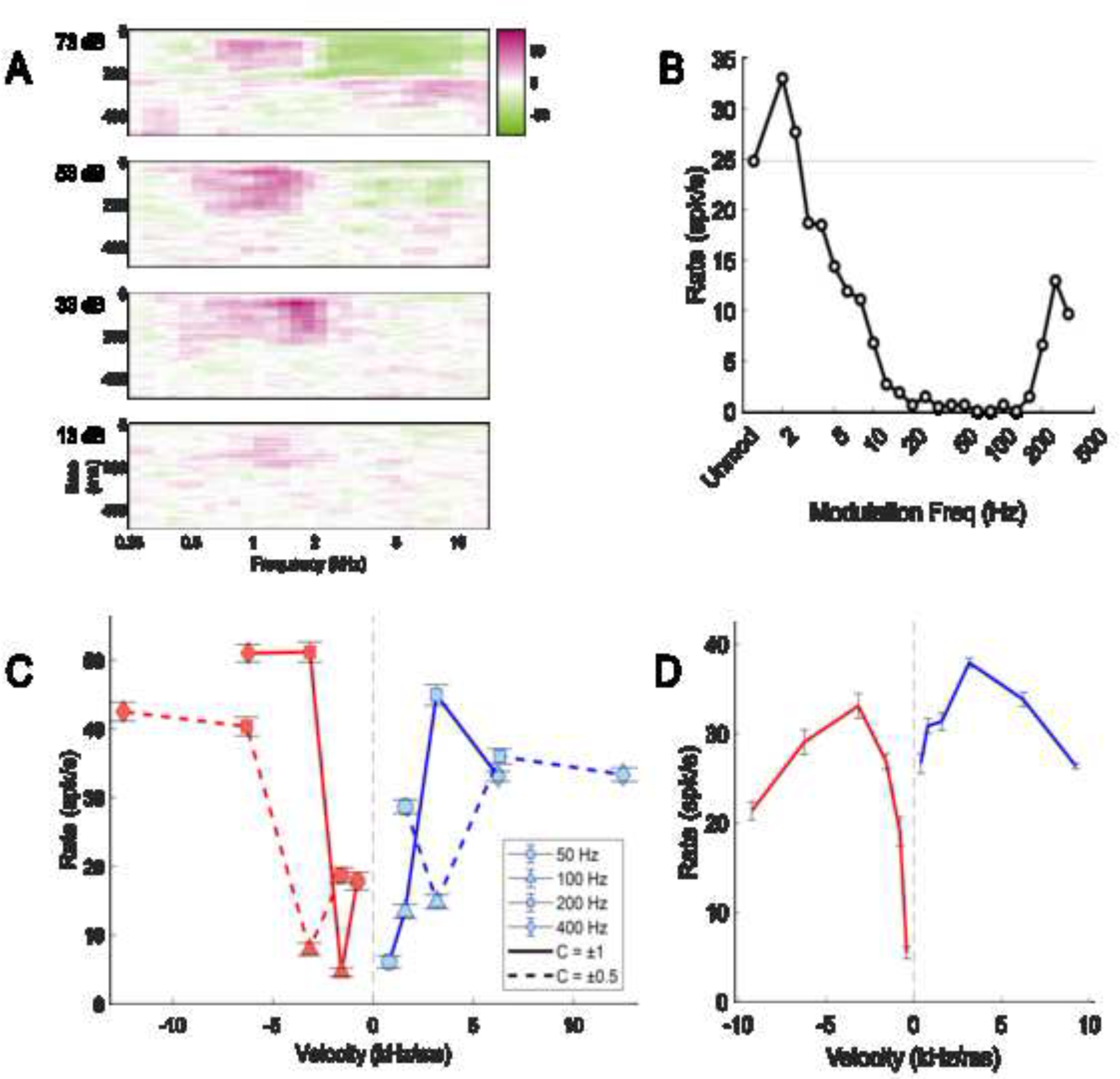
Responses of unit R24TT4P7N2 to characterizing stimuli. (A) Response Map, (B) Modulation Transfer Function, (C) SCHR dot-raster plot, (D) Rate-Velocity Function. Figure description identical to Fig. 3.

### 3.2. The prevalence of direction selectivity is influenced by stimulus periodicity

The prevalence of direction selectivity in response to both SCHR and aperiodic-chirp stimuli was quantified for the population of recorded neurons using ROC analysis. SCHR response rates were organized by F0 and C magnitude and grouped by chirp direction (sign of C). Aperiodic-chirp responses were compared between chirps of equal speed and opposite direction. Neurons for which response rate allowed direction to be discriminated 70.7% of the time via the ROC test were labeled direction-sensitive for the discriminated pair.

The ROC results across MTF groups suggest that prevalence of chirp-direction selectivity was impacted by the periodicity of the stimulus. When separated into groups based on MTF type, BS neurons were significantly less likely to be direction-selective to SCHR chirps than to aperiodic chirps (χ2 test of independence, Fig. 5). These significant differences were seen for F0s of 50 and 100 Hz, which correspond to typical modulation-frequency ranges for troughs in BS MTFs (Kim et al., 2020), and were likely a result of the SCHR-stimulus periodicity—the response to these SCHR F0s was suppressed, reducing sensitivity to chirp direction (as in Fig. 4). Furthermore, the prevalence of direction selectivity in BE and hybrid neurons was not significantly different between SCHR and aperiodic chirps for any chirp speed or SCHR F0 (Fig. 5). Responses to SCHR F0s that corresponded to peak modulation frequency values in BE and hybrid neurons were relatively unaffected, compared to responses to other SCHR F0s—for example, in Fig. 3C the direction bias at SCHR F0 50, 100, and 200 Hz was close to that of the corresponding velocities in the RVF (Fig 3D). However, the absence of direction bias at SCHR F0 400 Hz (Fig. 3C) demonstrates that chirp-direction selectivity may still be affected for an F0 outside the enhanced band. Overall, neurons were generally more sensitive to aperiodic chirps than to SCHR chirps, suggesting that stimulus periodicity may partially suppress chirp-velocity sensitivity in SCHR responses, especially in BS neurons.

**Figure 5.**
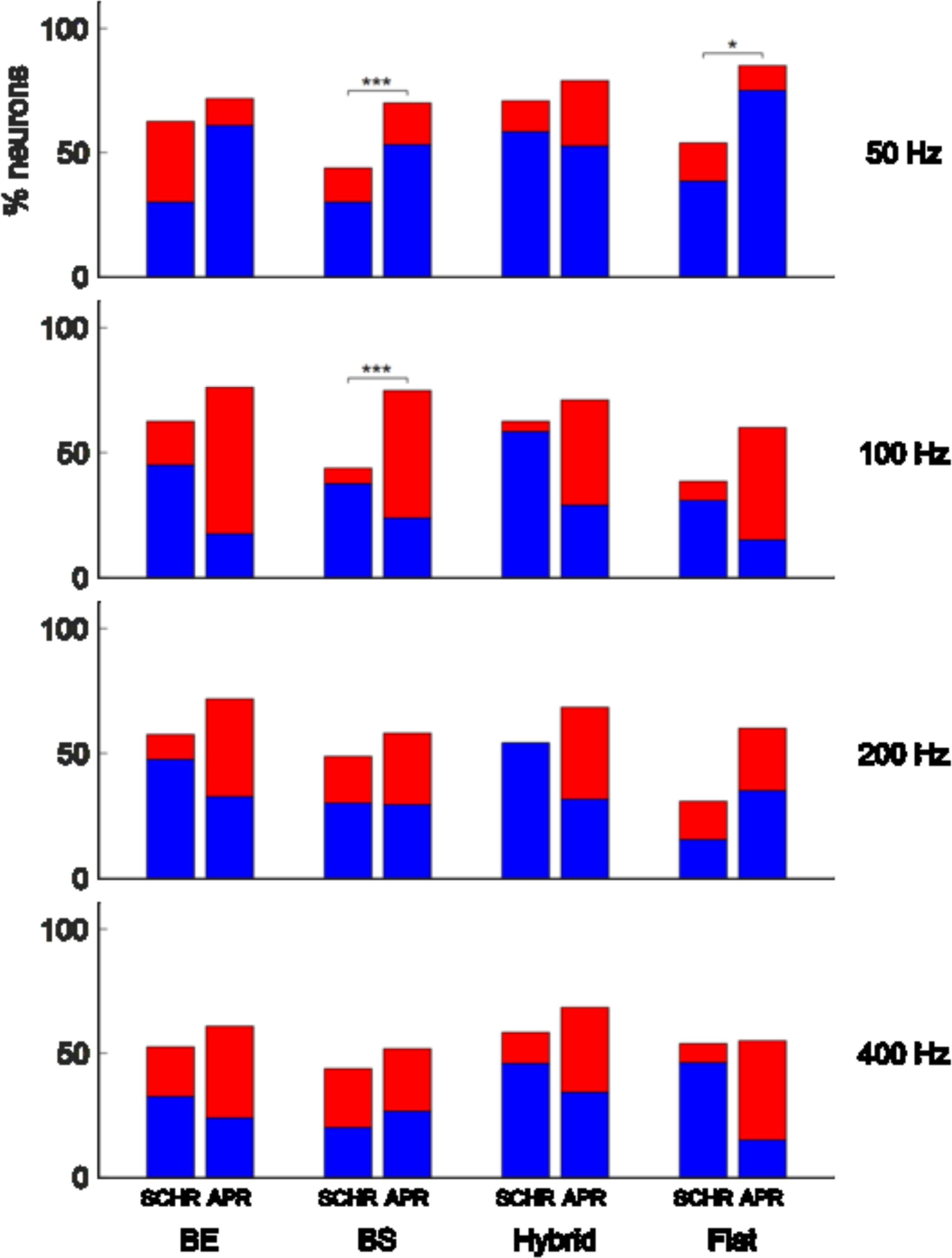
Percent of direction-selective neurons determined by ROC analysis for different MTF shapes. For each MTF shape, the left bar shows the percent of neurons selective for SCHR chirps, and the right bar shows the percent selective for aperiodic chirps (labeled APR). Chart row reflects the stimulus F0 for SCHR selectivity, or the equivalent SCHR F0 for aperiodic chirp-direction selectivity (the corresponding absolute velocities are 0.80, 1.59, 3.16, and 6.24 kHz/ms). Blue bar-segments correspond to upward-selective neurons, and red bar-segments correspond to downward-selective neurons. Brackets indicate significant differences of direction selectivity prevalence between groups, as determined by the χ^2^ test of independence, with asterisks indicating the level of significance (* p < 0.05, *** p < 0.001). p-values for each significant difference were 50 Hz—BS, p = 0.0001; 50 Hz—Flat, p = 0.0496; 100 Hz—BS, p < 0.0001.

Separating the ROC results by CF-range shows that stimulus periodicity impacts the prevalence of chirp-direction selectivity especially for higher-CF neurons (Fig. 6). Neurons in the high-CF (>6 kHz) range were direction-selective significantly more often for aperiodic chirps than for SCHR chirps, compared to low-CF (<3 kHz) and mid-CF (3-6 kHz) neurons (χ2 test of independence, Fig. 6 caption). This trend in high-CF neurons was observed for 50-, 100-, and 200-Hz-equivalent chirps, but not for 400-Hz-equivalent chirps.

**Figure 6.**
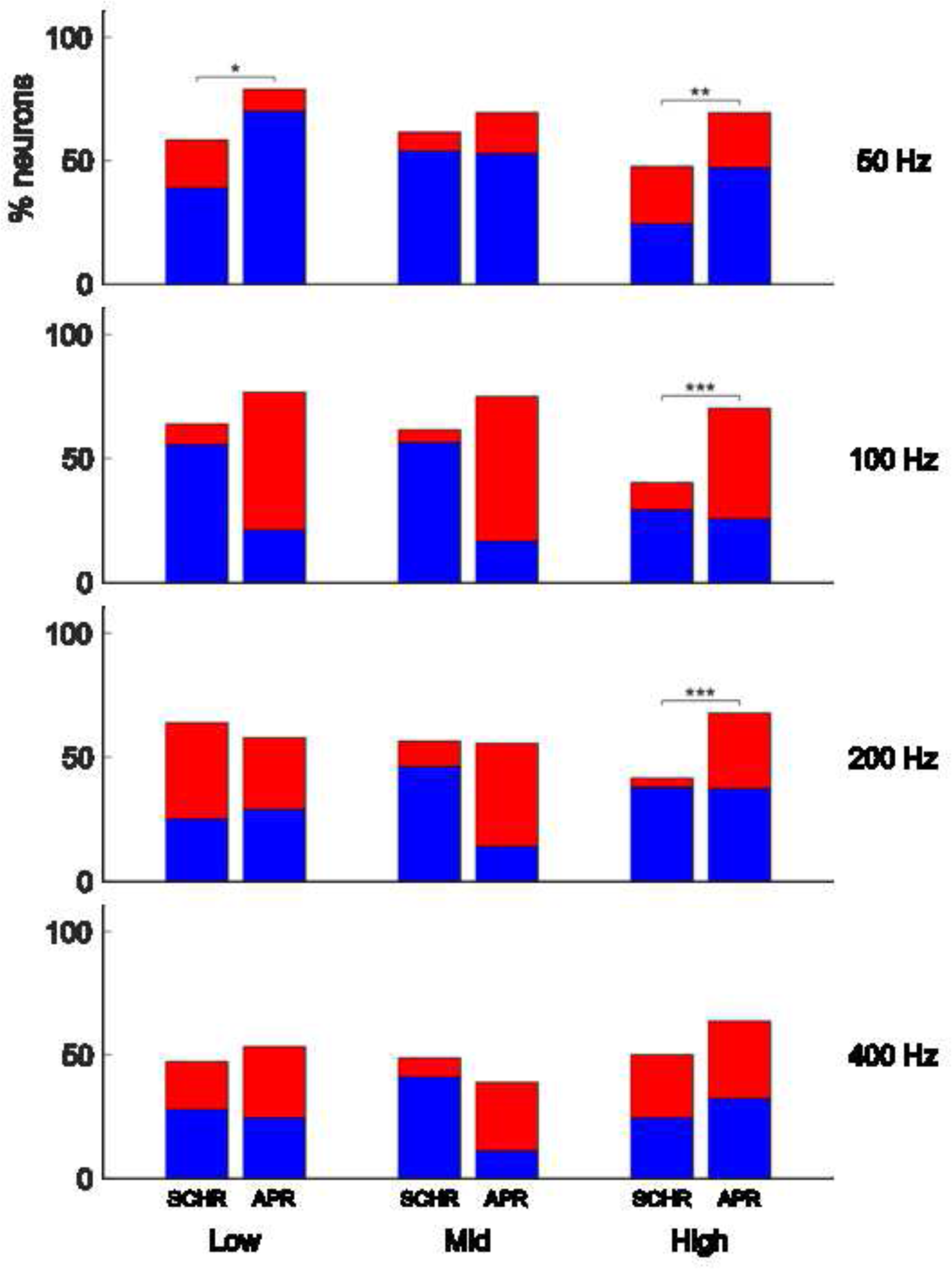
Percent of direction-selective neurons determined by ROC analysis for different CF groups—low (<3 kHz), mid (3-6 kHz), and high (>6 kHz). For each CF group, the left bar shows the percent of neurons selective for SCHR chirps, and the right bar shows the percent selective for aperiodic chirps (labeled APR). Chart row reflects the stimulus F0 for SCHR selectivity, or the equivalent SCHR F0 for aperiodic chirp-direction selectivity (the corresponding absolute velocities are 0.80, 1.59, 3.16, and 6.24 kHz/ms). Blue bar-segments correspond to upward-selective neurons, and red bar-segments correspond to downward-selective neurons. Brackets indicate significant differences of direction selectivity prevalence between groups, as determined by the χ^2^ test of independence, with asterisks indicating the level of significance (* p < 0.05, ** p < 0.01, *** p < 0.001). p-values for each significant difference were 50 Hz—Low, p = 0.0189; 50 Hz—High, p = 0.0018; 100 Hz—High, p < 0.0001; 200 Hz—High, p = 0.0002.

To quantify the prevalence of chirp-selectivity, the number of units that were selective at a minimum of one condition was determined. For SCHR responses, 90.5% of neurons had at least one direction-selective F0/C pair. For responses to aperiodic chirps, 99.6% of neurons had at least one direction-selective velocity pair. To ensure that a reasonable threshold was used for determining direction selectivity, responses to each chirp-direction pair were shuffled for each neuron, and an ROC test was performed on the shuffled responses. After shuffling the responses 1000 times, 3.2% of the neurons were selective at a minimum of one SCHR F0/C condition, and 13.3% were selective at a minimum of one aperiodic-chirp velocity pair. It is clear that the high prevalence of chirp-direction selectivity could not be due to variance in the data, suggesting that the criterion used in the ROC test (70.07%) was sufficiently high.

### 3.3. Periodicity sensitivity has stronger influence on SCHR rate responses than does chirp-velocity sensitivity

We quantified the dependence of responses to SCHR stimuli on both AM (periodicity) tuning and velocity sensitivity. Using a GLM, MTFs and RVFs were used to predict SCHR responses. Figure 7 shows a visualization of this strategy for an example neuron (the same neuron depicted in Fig. 3). The SCHR response is shown as a surface plot on periodicity-velocity axes (Fig. 7A); darker colors correspond to higher response rates. This neuron responded more strongly to lower F0s and to downward SCHR. The shaded bars above and to the left of the SCHR surface are the MTF and RVF, respectively, and describe how the response rate depended on periodicity or velocity alone. Using a weighted combination of these two responses, the GLM predicted the SCHR response (Fig. 7B). By fitting an equation that included both periodicity and velocity terms (Eqn. 1), an accurate prediction of the recorded SCHR data was made (*R*^2^ = 0.98). Removing either the periodicity or velocity terms reduced the quality of the prediction (Fig. 7C, D). Predicting the SCHR response using the periodicity-only model (Eqn. 2) resulted in the disappearance of the distinct direction selectivity at lower F0s (Fig. 7C). As expected, a model using only periodicity tuning could not explain the observed direction selectivity. Conversely, a prediction of the SCHR responses using the velocity-only model (Eqn. 3) explained a relatively small amount of variance in the data (*R*^2^ = 0.30). For most neurons (95%), more variance in the data was explained by the periodicity-only model than by the velocity-only model.

**Figure 7.**
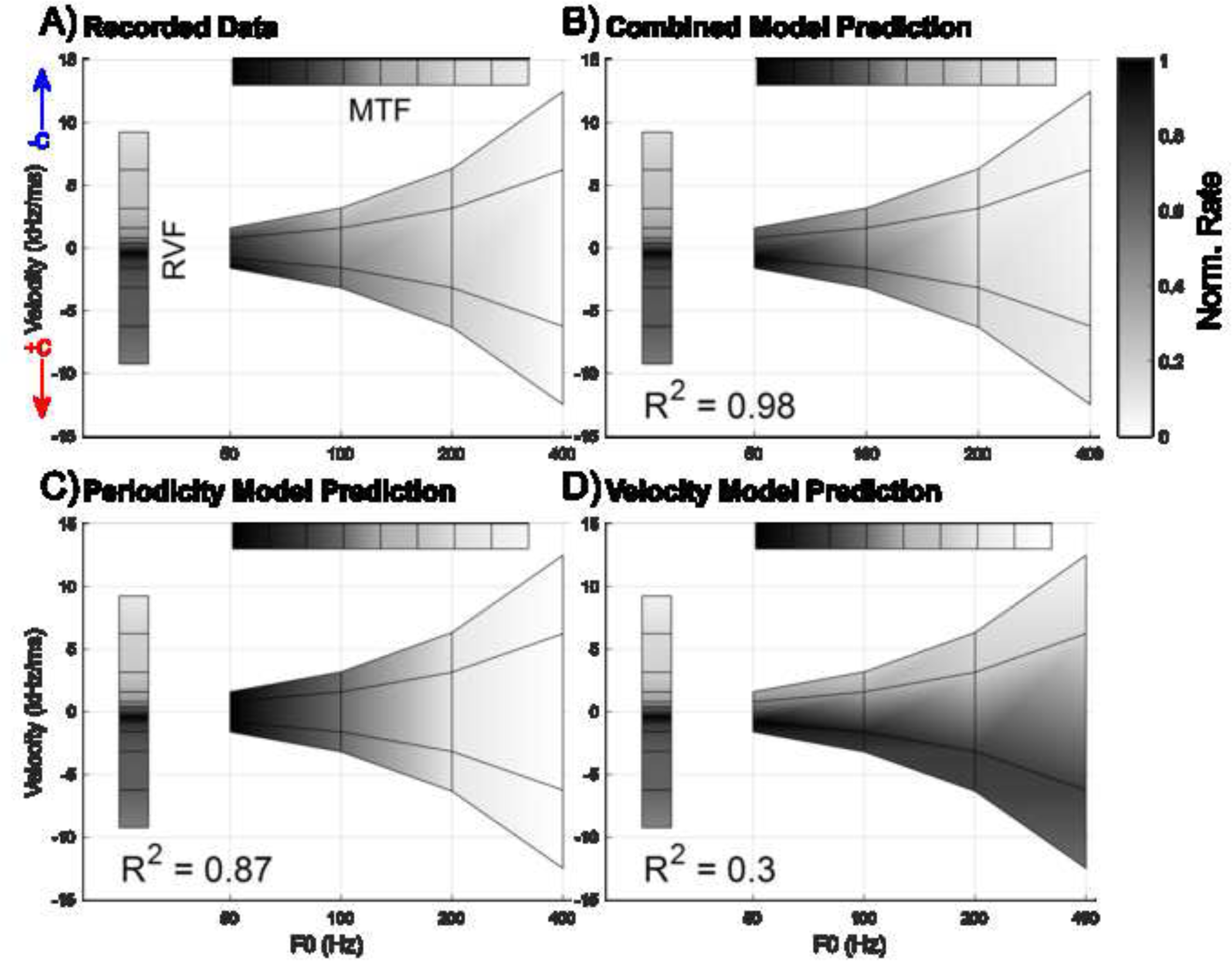
GLM prediction of SCHR rates for an example neuron (R24TT2P8N4; the same neuron as in Fig. 3). (A) SCHR rates plotted on periodicity-velocity axes (center surface). Each of the 16 points on the surface plot correspond to an individual SCHR F0/C-value combination – for instance, four points have a SCHR F0 of 50 Hz. Their velocities vary with C-value; the points’ C-values, from top to bottom, are -0.5, -1, 1, 0.5. This surface plot demonstrates the effect of C-value on chirp velocity. The periodicity and velocity responses are plotted alongside the SCHR surface plot—these are the MTF and RVF, respectively. The modulation frequency range of the MTF was limited to the section relevant to the range of SCHR F0s (50 – 400 Hz). Rate is marked using color, with darker shades indicating larger rates. The rate scale of each surface plot is separately normalized by its own maximum rate—thus, shading does not reflect absolute rates. Surface plot colors are interpolated. (B) SCHR rates predicted using the combined GLM equation (Eqn. 1), which uses both periodicity and velocity cues. Coefficient of determination (R^2^) between predicted and recorded SCHR rates is also shown. (C) SCHR rates predicted using the periodicity-only GLM equation (Eqn. 2), and corresponding R^2^-value. (D) SCHR rates predicted using the velocity-only GLM equation (Eqn. 3), and corresponding R^2^-value.

Figure 8 illustrates the interaction of stimulus periodicity and velocity in the combined GLM model. For aperiodic-chirp velocities that closely matched the velocities of SCHR chirps with F0 equal to 50 Hz, a large aperiodic-chirp response rate predicts a large SCHR response rate (blue line). For higher F0s, aperiodic-chirp response rate had a smaller impact on the predicted SCHR response rate, with slopes approaching zero (200- and 400-Hz lines, yellow and purple). Therefore, despite the RVF suggesting a strong selectivity for downward sweeps at all velocities, Fig. 8 shows that the interaction between velocity and periodicity sensitivities resulted in lower SCHR response rates for high F0s and suppressed direction selectivity. Similar velocity-periodicity interactions as illustrated in Fig. 8 were typical of many neurons in the study; the interaction term, *b*_*I*_, for 24% of neurons was statistically significant in predicting SCHR response rates (two-sided t-test, p < 0.05). In comparison, the periodicity coefficient, *b*_*P*_, was statistically significant for 40% of neurons, and the velocity coefficient, *b*_*V*_, was significant for 7%.

**Figure 8.**
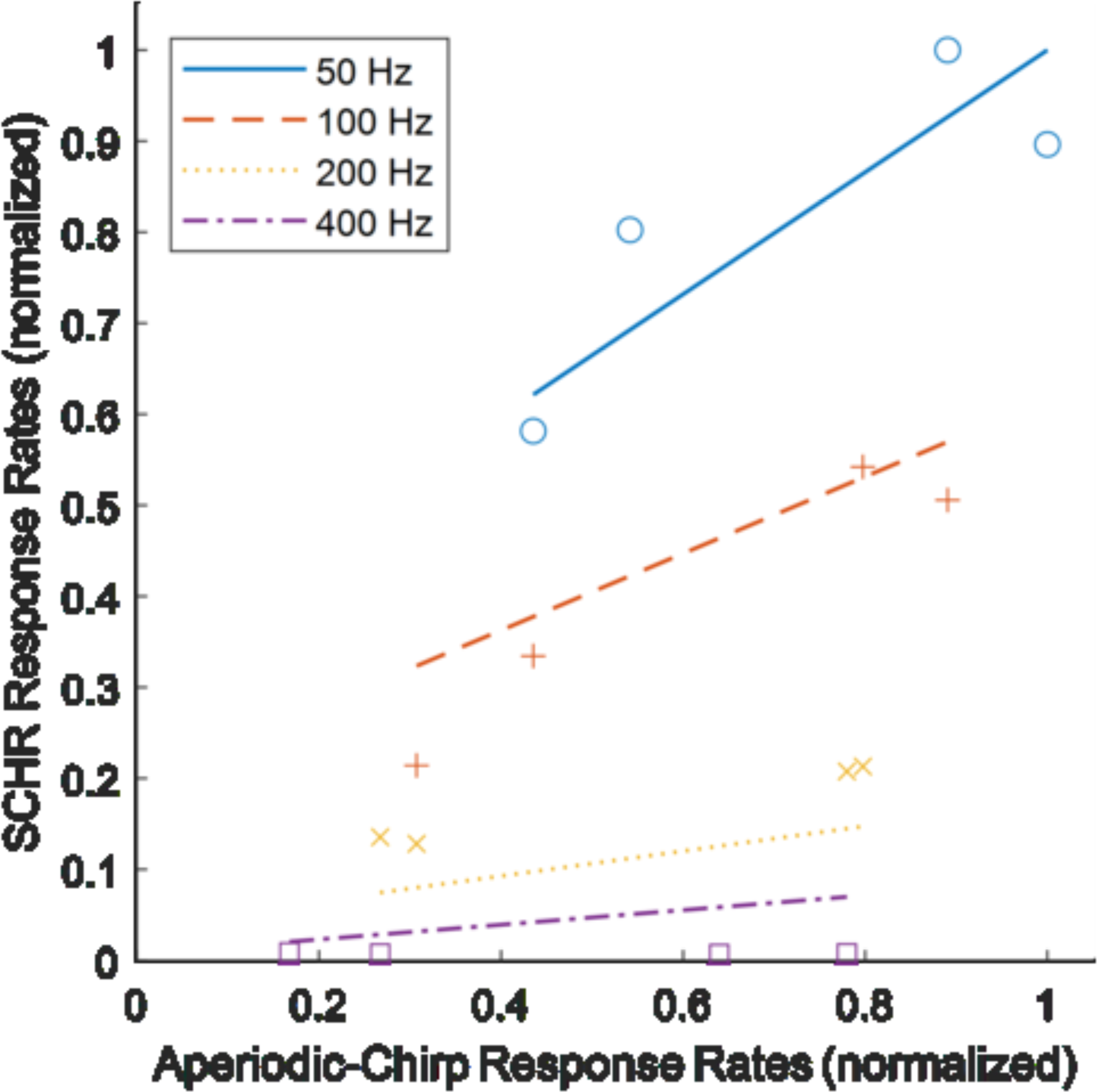
SCHR response rates vs aperiodic-chirp response rates of equivalent velocity for R24TT2P8N4. GLM predictions are shown as lines. Blue solid line and circles correspond to 50 Hz, orange dashed line and plus signs correspond to 100 Hz, yellow dotted line and crosses correspond to 200 Hz, and purple dash-dotted line and squares correspond to 400 Hz. Both SCHR and aperiodic-chirp rates are normalized by their own maximum rate. This example neuron is also shown in Figs. 3 and 7.

Using the method outlined above (Fig. 7), an *R*^2^ value was obtained for each of the three GLM models (Eqns. 2-4) by comparing model predictions to neural responses. The contribution of the velocity term to the total explainable variance was calculated as a percentage, using Eqn. 4. Figure 9 shows this percentage versus the variance explained by the combined model. For 95% of the neurons studied, periodicity was a more important feature than velocity in predicting SCHR response rates. In neurons with high explainable variance (80-100%), the contribution of velocity rarely exceeded 20%. Velocity was usually more important in neurons with a low overall explainable variance. These results demonstrate that, while velocity sensitivity is critical to explain chirp-direction sensitivity, the periodicity tuning tends to dominate the SCHR response rates.

**Figure 9.**
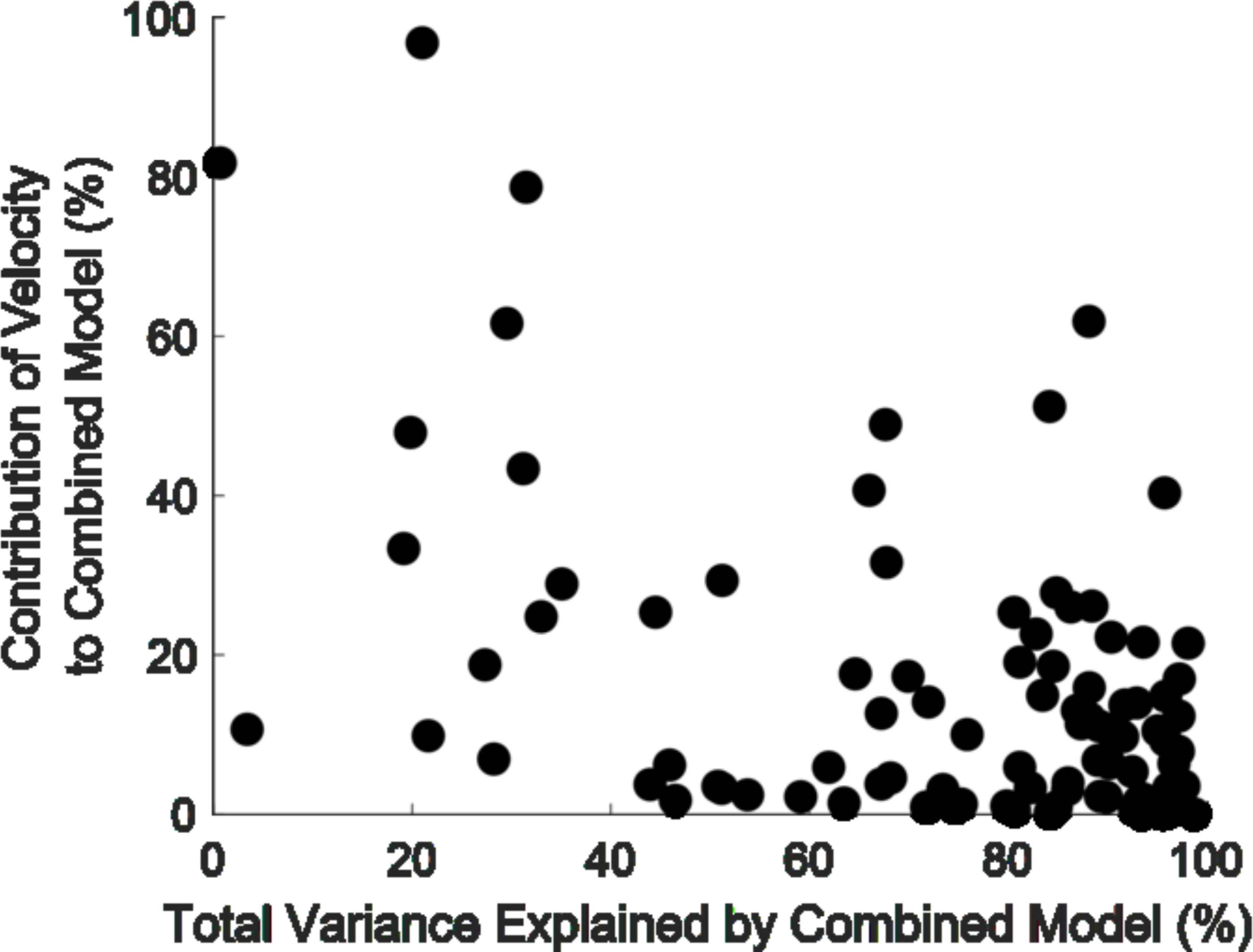
Percent contribution of the velocity terms to the total explainable variance by the combined model vs. variance explained by the combined model.

### 3.4. RVFs were often direction-selective at slower velocities and direction-insensitive at higher velocities

Responses to aperiodic chirps isolate neural sensitivity to chirp velocity, so it is interesting to consider trends in these responses across cell groups. Figure 10 shows the results of ROC analysis for aperiodic-chirp responses, for neurons categorized by MTF type (Fig. 10A) and CF range (Fig. 10B). For both MTF type or CF range, there were no systematic differences between groups in prevalence of chirp-direction selectivity. Additionally, direction bias between both MTF groups and CF groups for a given chirp speed were comparable. However, one notable trend was that prevalence of selectivity decreased with increasing chirp speeds. Additionally, upward-selectivity was more common for lower-speed chirps (0.80 kHz/ms absolute velocity and below), and downward-selectivity was more common for higher-speed chirps (1.59 kHz/ms absolute velocity and above). Overall, for aperiodic-chirp responses, there was not a trend across MTF types or CF, in either prevalence of direction selectivity or direction of bias. The direction selectivity observed was diverse, with both upward- and downward-selective units seen for every category.

**Figure 10.**
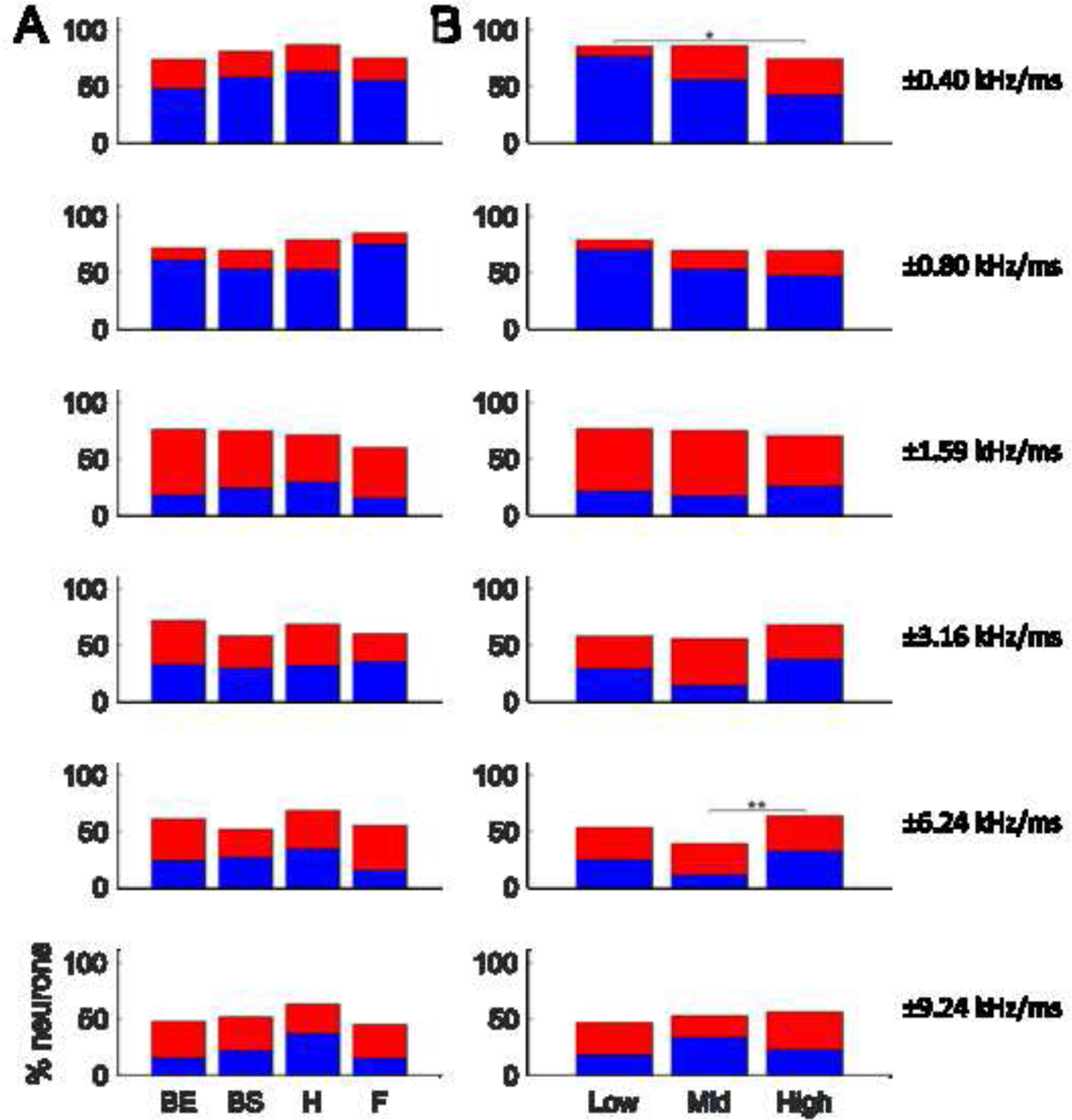
Percent of direction-selective neurons determined by ROC analysis, by aperiodic-chirp velocity. (A) Neurons are grouped by MTF shape (BE – Band-Enhanced, BS – Band-Suppressed, H – Hybrid, F – Flat). (B) Neurons are grouped by CF range—low (<3 kHz), mid (3-6 kHz), and high (>6 kHz). Rows correspond to velocities of chirp duration, with velocities ±0.40, ±0.80, ±1.59, ±3.16, ±6.24, and ±9.24 kHz/ms corresponding to equivalent SCHR F0s of 25, 50, 100, 200, 400, and 600 Hz, respectively. Blue bar-segments correspond to upward-selective neurons, and red bar-segments correspond to downward-selective neurons. Brackets indicate significant differences of direction selectivity prevalence between groups, as determined by the χ^2^ test of independence, with asterisks indicating the level of significance (* p < 0.05, ** p < 0.01). p-values for each significant difference were ±0.40 kHz/ms—Low-High, p = 0.0481; ±6.24 kHz/ms—Mid-High, p = 0.0083.

As demonstrated by the ROC analysis performed on aperiodic-chirp responses, significant direction selectivity to aperiodic chirps was commonplace. However, this analysis only addressed selectivity between direction of chirp-pairs and not sensitivity to chirp velocity. Although RVFs revealed velocity sensitivity to aperiodic chirps for individual neurons (Figs. 3-4), further analysis was needed to characterize trends in velocity sensitivity for the population of neurons. In order to identify the most common RVF characteristics, principal component analysis (PCA) was performed on the set of RVFs obtained from the population of single-units (247 in total). This approach reduced the dimensionality of the RVF dataset and extracted the principal components, or the features that explained the most variability in the population.

Figure 11 shows the first three principal components, which together explained 74.0% of the variance in the RVFs. Principal component 1 (PC1) had relatively high weights across high-magnitude velocities (3.16 kHz/ms and faster), showing that rate responses to high-velocity chirps tended to be positively correlated, regardless of direction. In other words, neurons responded similarly to either direction of high-velocity chirp. PC1 explained 48.5% of variance in the RVF data. Principal component 2 (PC2) had high weights at low-magnitude velocities (1.59 kHz/ms and slower), showing that rate responses to low-velocity chirps also tended to be positively correlated, regardless of direction. PC2 explained 13.8% of variance in the data. Finally, PC3 had large positive weights at slow upward velocities and negative weights at slow downward velocities (1.59 kHz/ms and slower), demonstrating that responses to upward and downward velocity chirps tended to be negatively correlated at slow velocities. PC3 was the main PCA dimension reflecting direction selectivity. PC3 explained 11.7% of variance in the data.

**Figure 11.**
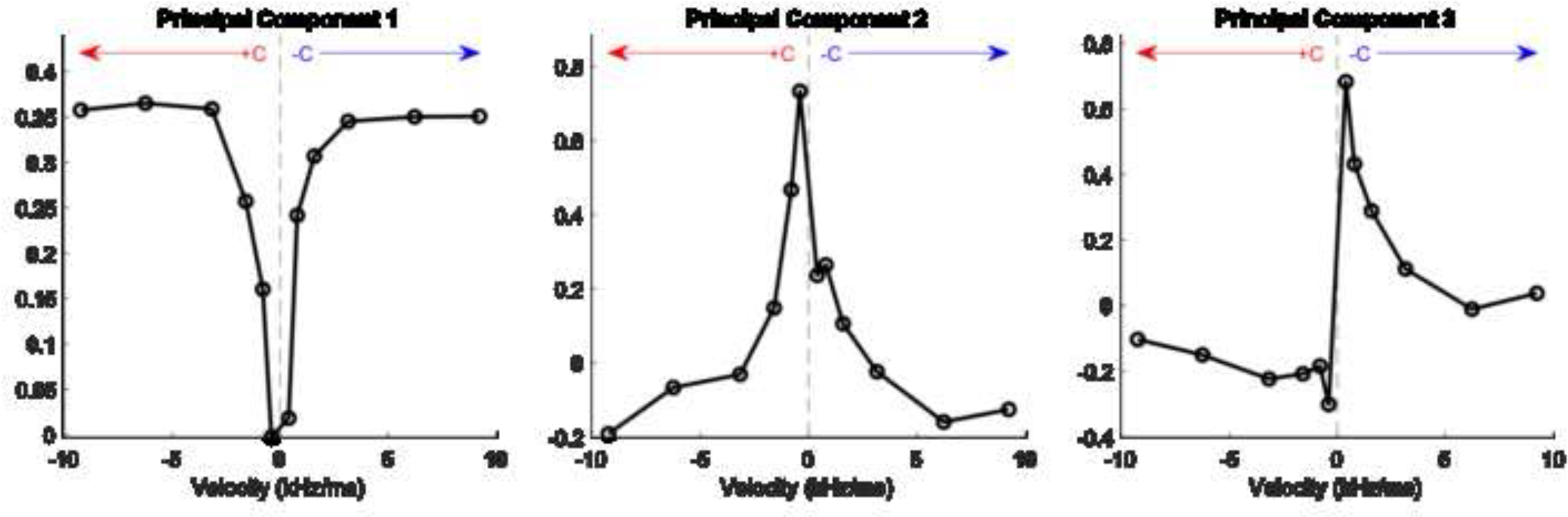
Principal components derived from the set of RVFs (left, PC1; center, PC2; right, PC3). Y-axes reflect principal-component weights and are unitless.

The shapes of the first two principal components of the RVF data suggest that the most salient features of the RVF were the responses to high- and low-velocity chirps, rather than the direction of those chirps, with the boundary between these groups located at about 2 kHz/ms. PC3 showed that the direction selectivity was an important feature of these datasets; specifically, PC3 primarily represented selectivity to the direction of low-velocity chirps. In contrast, neural responses to chirps of high velocities depended more on chirp speed (magnitude of velocity) rather than direction, with the divide between low- and high-velocity chirps at about 2 kHz/ms. These results are consistent with the observation that direction selectivity to high-velocity chirps was not as common (Fig. 10).

Principal-component scores provide a measure of how well each RVF aligns with the principal components in Fig. 11, and thus are a useful way to divide the population based on type of velocity sensitivity. This approach also allowed a comparison of velocity sensitivity against other response properties, such as MTF type and CF. As shown in Fig. 12, there was not a clear relationship between the scores of principal components 1-3 and either MTF type (symbol shapes) or CF (symbol colors). Furthermore, based on the scatter plots in Fig. 12, there were no clusters of neurons based on principal-component analysis – that is, RVF subgroups were not revealed by this analysis. This result suggests that a neuron’s RVF was not directly related to its MTF or CF.

**Figure 12.**
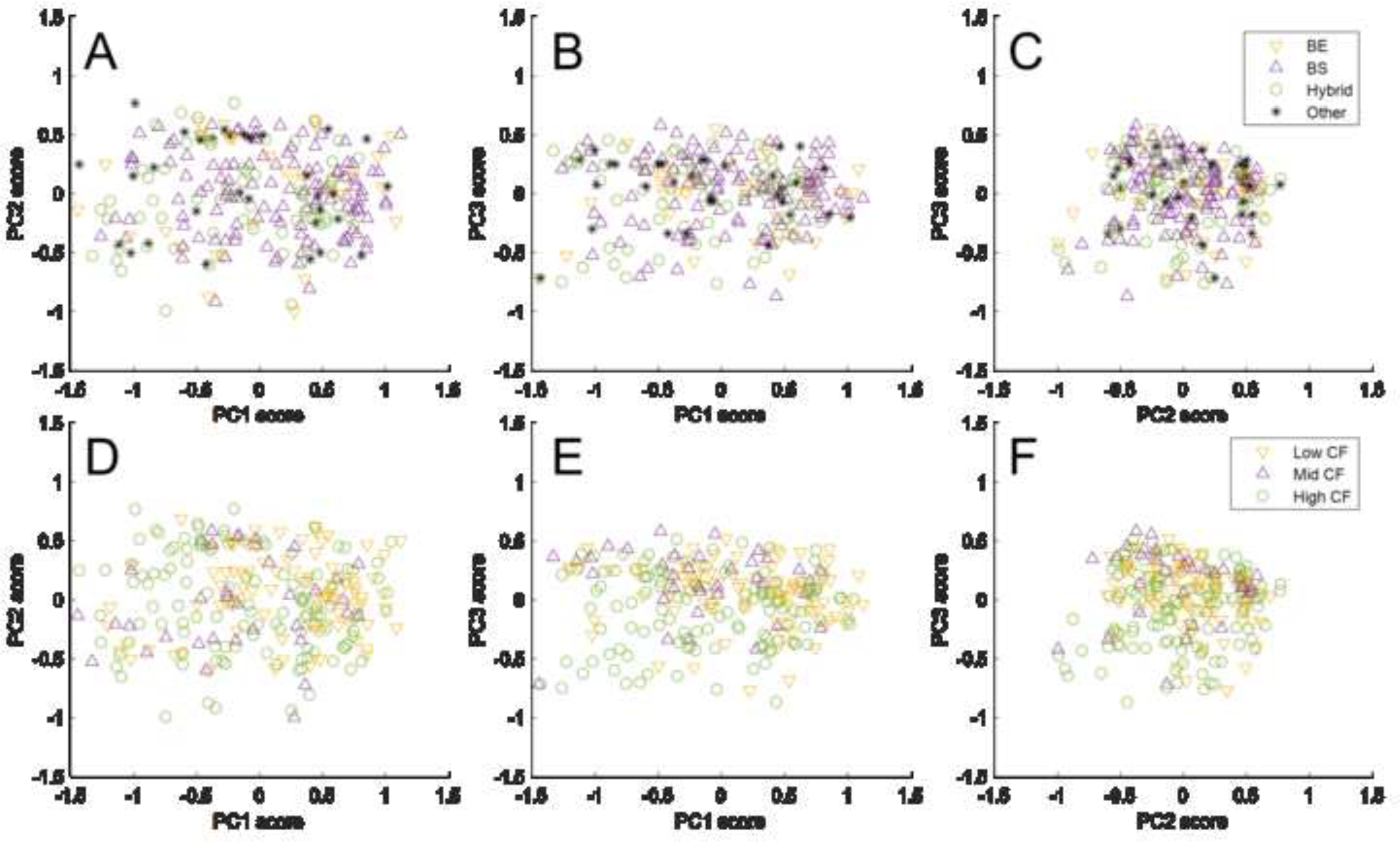
Scatter plots depicting the Principal Component scores of each neuron, organized by MTF shape (A-C) and CF range (D-F). (A and D, PC2 vs PC1; B and E, PC3 vs PC1; C and F, PC3 vs PC2). In the top row (A-C), BE neurons are represented by yellow up-pointing triangles, BS neurons by purple down-pointing triangles, hybrid neurons by green circles, and other MTF shapes by black asterisks. In the bottom row (D-F), Low-CF neurons are represented by yellow up-pointing triangles, mid-CF by purple down-pointing triangles, and high-CF by green circles.

## 4. Discussion

This study demonstrated the diversity of sensitivity to the velocity of fast frequency chirps in the IC, for the first time in an awake rabbit model. Additionally, this study introduced aperiodic-chirp stimuli as a way to isolate neural velocity sensitivity from periodicity tuning. ROC analysis was used to calculate the prevalence of chirp-direction selectivity for responses to SCHR and aperiodic chirps. BS neurons were direction-selective significantly less often for SCHR chirps than for aperiodic chirps, suggesting that chirp sensitivity was suppressed for SCHR stimuli with F0s in the trough of BS MTFs. For all MTF-types, periodicity tuning influenced the overall rate responses to SCHR stimuli. Additionally, GLM analysis showed that more variance in SCHR data could be explained by periodicity than by velocity. This study also showed that individual neurons had a wide variety of rate-response profiles to aperiodic chirps, described using RVFs. RVFs revealed that neurons were more commonly direction-selective to chirps with velocities below 2 kHz/ms than above, and responded to chirps of velocities greater than 2 kHz/ms in a more direction-insensitive way.

A common use of Schroeder-stimuli has been as a masker in psychophysical experiments evaluating detection of an increment in tone level. Notably, there is an impact of Schroeder-phase on the effectiveness of SCHR stimuli as maskers, with up to 20-dB lower thresholds (less masking) for downward-SCHR maskers than for upward-SCHR maskers (Smith et al., 1986). An explanation from the literature attributes this effect to asymmetry of filtering in the cochlea: phase-dispersive properties of the basilar-membrane (BM) result in a peakier temporal response to downward than for upward SCHR stimuli. The peakier response is then compressed at the output of the cochlea, resulting in higher average energy in upward-SCHR BM responses compared to downward-SCHR responses, and thus more effective masking by upward SCHR stimuli (Recio and Rhode, 2000; Summers et al., 2003). This logic can also be applied to explain rate-differences between SCHR-chirp directions. However, the results here suggest that other factors underlie the widespread chirp-direction selectivity of IC single-units observed in this study. First, the phase-dispersion of the BM depends on CF (Ruggero et al., 1997; Carney et al., 1999; Shera, 2000), with neural masking differences in chinchilla observed primarily for CFs > 3-4 kHz (Recio, 2001). In this study, no consistent differences were observed in either the prevalence or direction of chirp-direction selectivity between low-CF (< 3 kHz) and mid- or high-CF (> 3 kHz) neurons in aperiodic-chirp response rates (Fig. 10b). Secondly, the cochlear dispersion hypothesis can only explain selectivity towards upward chirps; despite this, neurons with CF > 3 kHz could be selective for either direction of chirp (Fig. 10b), suggesting that cochlear phase-dispersion was not the basis of IC chirp sensitivity. Similarly, Henry et al. (2023) found diverse SCHR selectivity in the IC of budgerigars (parakeet), though a statistically significant gradient was observed from upward SCHR selectivity at CFs below 2-3 kHz to downward SCHR selectivity for higher CFs. These findings are also consistent with a recent study in gerbils (Steenken et al., 2022) that reported that average rates of AN fibers could not explain behavioral discrimination of SCHR stimuli, whereas temporal analyses of AN responses over short (1-ms) time windows could explain behavioral thresholds. Low correlations of AN rate to behavior provide further evidence against cochlear phase-dispersion driving IC selectivity. On the other hand, the presence of information for SCHR-direction discrimination in the relatively fine timing of AN fibers is consistent with neural mechanisms that combine temporal information across frequency channels. Finally, a recent study has shown that phase curvature of the extreme apical region of guinea pig and gerbil cochleae is different from other regions (Recio-Spinoso et al., 2023), complicating the statement that downward chirps always have peakier BM responses. This unique phase curvature may influence processing of chirps containing low-frequency components, such as the SCHR and aperiodic chirps used in this study.

A potential mechanism to explain the diverse chirp-response profiles seen in this study may be inhibitory inputs to the IC. IC neurons receive excitatory and inhibitory inputs with varying frequencies and latencies that may interact to produce chirp-direction selectivity (Pollak et al., 2011). Interactions between excitatory and inhibitory inputs varying in latency have long been implicated in mechanisms for frequency-sweep selectivity in bats (Suga, 1965; Fuzessery and Hall, 1996; Gordon and O’Neill, 1998). These mechanisms are broadly dependent on sideband inhibitory regions in excitatory tuning curves and the resulting asymmetry of responses to upward and downward frequency sweeps—this inhibition may arise from a variety of IC inputs. While not all response maps showed inhibitory sidebands, it is interesting to examine those that did within the framework of this theory. For instance, the neuron in Fig. 4 had an above-CF inhibitory sideband, with a longer duration and shorter latency than the excitatory band at higher sound levels. The location of this inhibitory sideband would predict a stronger response to upward chirps. Indeed, the RVF of this neuron is mostly biased towards upward chirps. The contribution of off-CF inhibition observed in response maps to interpreting chirp-direction sensitivity in the IC population studied here was precluded by the low-spontaneous rates of most IC neurons. To more comprehensively assess the effect of inhibition on chirp sensitivity, response maps using pure tones in background noise could be measured— this would be an interesting direction for a future study.

Another mechanism for chirp-direction selectivity was proposed by Rall (1969), based on dendritic filtering of several excitatory inputs organized spatially by frequency. In Rall’s framework, a sweep that activates distal dendrites first, and proximal dendrites later, would temporally sum and generate a large synaptic input. This mechanism has been hypothesized to underlie the frequency-sweep direction selectivity of octopus cells in the posteroventral cochlear nucleus (Godfrey et al., 1975; Rhode and Smith, 1986; Lu et al., 2022). Notably, octopus cells project to the ventral nucleus of the lateral lemniscus (VNLL), which provides inhibitory innervation to the IC (Adams, 1997; Vater et al., 1997; Winer et al., 1995). Thus, direction selectivity of octopus cells, by way of the VNLL, may explain chirp-direction selectivity in the IC; in this scenario, responses to chirps in one direction would be inhibited by input from the VNLL and disinhibited by chirps in the opposite direction. The possibility that IC chirp sensitivity originates in octopus cells is particularly intriguing because, like IC neurons, octopus cells display a diversity of chirp response profiles and can be selective to either upward or downward chirps (Lu et al., 2022). Note that Lu et al. (2022) suggested that the diverse chirp-direction selectivity in octopus cells could be explained by sequence detection, with response magnitude determined by a sequence of inputs with both frequency-dependent delays and varied amplitudes, enabled by the relatively long hyperpolarization of low-threshold potassium channels. Another possibility is that chirp-direction selectivity may originate in stellate neurons of the cochlear nucleus, which have been noted to display strong-upward SCHR selectivity, at least for higher CFs (Recio and Rhode, 2000; Recio, 2001). However, because IC neurons can be selective for either upward or downward chirps (Fig. 10), cochlear nucleus stellates cannot be the only source of chirp sensitivity. Finally, another possibility is that chirp-direction selectivity may arise at the level of the IC, from an inhibitory interneuron with inputs extending across iso-frequency laminae, such as IC stellate neurons (Oliver, 1984). The literature suggests that such neurons may also have low-threshold potassium channels (Sivaramakrishnan and Oliver, 2001) and could therefore be selective to a sequence of frequency inputs in a manner similar to that proposed for octopus cells.

The aperiodic-chirp stimulus was composed of discrete, single-chirps, whereas the SCHR stimulus was a longer, sustained harmonic tone. It is possible that the presence of silent gaps might partially explain differences in responses to the two stimuli. In the GLM analysis, the combined GLM with both periodicity and velocity terms sometimes could not explain a high percentage of the variance (Fig. 9), suggesting that additional factors shaped SCHR response rates. Long-term rate modulation in response to a sustained stimulus may be one of these factors. This long-term effect may be attributable to the auditory efferent system, and cannot be captured in brief aperiodic-chirp responses. Similar long-term changes in rate have been reported in response to other complex sounds, representing a possible direction for future research (Farhadi et al., 2021). Furthermore, one could imagine a future study designing an aperiodic-chirp stimulus containing no silent gaps.

SCHR-chirp sensitivity in the IC suggests a mechanism that may influence midbrain-level encoding of speech. When speech sounds, or any animal vocalizations, are produced, resonances within the vocal tract act as second-order filters, introducing phase shifts between components of harmonic sounds (Klatt, 1980). In the same way that SCHR chirps are produced by the relative phases of harmonic components, vocalizations contain chirp-like features to which IC neurons may be similarly sensitive. IC neurons in this study were sensitive to SCHR chirps and directions over a frequency range relevant to human speech. Chirps extended over the frequency range of formants (200 – 3000 Hz for F1 and F2), and SCHR F0s were within the range of F0s in human speech (about 100 to 400 Hz) (Ladefoged and Johnson, 2014). Therefore, IC sensitivity to chirp velocity may influence midbrain coding of speech, music, and other complex harmonic sounds. Also, responses to these sounds may be affected by the interaction of periodicity tuning and chirp-velocity sensitivity observed in this study.

## Acknowledgements

We acknowledge the assistance of Kristina S. Abrams, Johanna B. Fritzinger, and Dr. Swapna Agarwalla for help with physiological experiments. Funding: This work was supported by the National Institutes of Health [grant numbers NIH-DC001641 and NIH-F31DC019816].

## Author Statement

NA

## Notes

### Competing Interest Statement

The authors have declared no competing interest.

### Summary of Updates

Minor revisions based on feedback from reviewers.

